# The loss of ATRX/DAXX complex disturbs rDNA heterochromatinization and promotes development of glioma

**DOI:** 10.1101/745307

**Authors:** XiangRong Cheng, Qi Jiang, XingLin Hu, XingWei Huang, Liu Hui, YanJun Wei, Na Li, Nan Wang, JingLing Shen, Yan Zhang, Lei Lei

## Abstract

**Background:** Ribosomal DNA (rDNA) transcription by the RNA polymerase I (Pol I) is a rate-limited step for ribosome synthesis, which is critical for cell growth, cell differentiation, and tumorigenesis. Meanwhile rDNA transcription is modulated by DNA methylation and histone epigenetic modification. Though with great progress in epigenetic research recently, it still remains much uncertain about the relationship of histone variant epigenetic modification and rDNA transcription.

**Results:** In this study, epigenetic profiles of silent rDNA in next-generation sequencing datasets were examined. We found that the chaperone of histone variant H3.3, the alpha-thalassemia/mental retardation X-linked syndrome protein (ATRX)/death domain-associated protein (DAXX) complex, and methyltransferase SET domain bifurcated 1 (Setdb1, also known as ESET) help maintain H3.3K9me3 modifications among the promoter and coding regions of silent rDNA. Our experiments further confirmed that DAXX depletion leads to the conversion of silent rDNA into upstream binding factor-bound active rDNA and the release of rDNA transcriptional potency. Support for this model is provided by data from a low-grade glioma in which ATRX is lost and a higher level of ribosomal biosynthesis, nucleolus activity, and proliferation are observed.

**Conclusions:** We demonstrate a model of epigenetic regulation for rDNA with roles for the ATRX/DAXX complex and H3.3/H3K9me3 modifications identified. Thus, loss of ATRX/DAXX may represent a driving force for tumorigenesis due to its contribution to the release of rDNA transcriptional potency.

## INTRODUCTION

The transcriptional activity of ribosomal DNA (rDNA) arrays limits the rate of cellular ribosomal biogenesis (1). Mouse 47S rDNA arrays include 400 copies of tandem repeat genes, which are distributed among six chromosomes (12, 15, 16, 17, 18, and 19). In each repeat unit, the gene coding region and intergenic spacer (IGS) region constitute 45 kilobase pairs (kbp) and they exhibit very different biological functions. For example, promoter and coding (P&C) regions are characterized by a high GC content and are often responsible for the transcriptional activity of rDNA (2). Meanwhile, IGS regions tend to have normal GC content and are involved in mediating nucleolar matrix-attachments (3). Due to the presence of multiple copies of tandem repeat genes, there are two strategies which have been proposed for the regulation of rDNA transcription in cells: 1) growth factors, nutrients, or a range of stress factors regulate the transcriptional efficiency of RNA polymerase I (4), or 2) epigenetic modifications regulate the ratio of active rDNA to silent rDNA and transcriptional potency (5). The former strategy represents a short-term regulatory pathway, with nutrients, growth factors, and/or carcinogenic agents upregulate transcriptional activity, while senescence, toxic lesions, viral infections, and/or genotoxic or metabolic stresses downregulate transcriptional activity. Meanwhile, the latter strategy represents a long-term stable mechanism which would potentially play a critical role in differentiation and transformation via enhancement or inhibition of transcriptional potency. For differentiated cells which have a lower metabolic demand, it is necessary to reduce the number of copies of active rDNA and establish more silent rDNA in order to stabilize the level of transcription and establish heterochromatin (6). Conversely, certain cells undergo rapid proliferation and experience a huge demand for protein synthesis.

These cells include tumor cells (7), growing oocytes, and reprogrammed cells (8). Indeed, ribosomal biogenesis activity has been found to be directly related to cell proliferation, and doubling of a cell’s proliferation rate requires a four-fold increase in the rate of ribosomal biogenesis (9). Consequently, in rapidly proliferating cells, there is a critical need to convert silent rDNA into more active rDNA to boost transcriptional activity.

It is well known that silent rDNA is established by the nucleolar chromatin remodeling complex (NoRC) which consists of ATPase SNF2h and the nucleolar protein, transcription termination factor 1 (TTF1)-interacting protein 5 (TIP5). Nucleolar retention of NoRC requires interactions between TIP5 and long non-coding RNAs that originate from IGS DNA, referred to as pRNA (10). NoRC is then able to target rDNA promoters and establish heterochromatin by interacting with histone methyltransferases, histone deacetylases, poly (ADP-ribose) polymerase 1 (PARP1) (11), and DNA methyltransferases (DNMTs) to mediate the introduction of H3K9me2 and H4K20me3 modifications, deacetylate H4, and mediate the protein poly ADP-ribosylation (PARylation) and *de novo* DNA methylation of silent rDNA (11,12). However, neither knockdown of TIP5 (13), nor overexpression of pRNA (6), has been shown to affect H3K9me3 modifications on rDNA. Therefore, it remains unclear how H3K9me3 modifications are established on silent rDNA.

In recent studies of histone variant H3.3 in relation to epigenetic mechanisms in the genome (14–16), H3.3 has been shown to differ from canonical H3 by 4 or 5 amino acids and it is incorporated into chromatin independent of DNA replication (17). An important link between H3.3 and its chaperones, HIRA or alpha-thalassemia/mental retardation X-linked syndrome protein (ATRX)/death domain-associated protein (DAXX), has been explored (14,15,18). HIRA mediates H3.3 turnover mainly in association with active transcription (14), ATRX serves as a recruiter protein by recognizing H3K9me3 via its ADD domain (18), and DAXX acts as a chaperone via its N-terminus to interact directly with variant-specific residues of H3.3 (15). The ATRX/DAXX complex then deposits H3.3 and modifies the histone tail to establish H3.3K9me3 with assistance from the lysine methyltransferase, SET domain bifurcated 1 (Setdb1, also referred to as ESET) or Suv39h1/2. The latter has been reported to play a role in telomeres (14,15), pericentric heterochromatin (19), and other sites (20). Despite the mechanism of ATRX/DAXX complex being well-understood, it remains unclear whether it is involved in epigenetic control of rDNA.

The relationship between ATRX and rDNA was first reported based on immunofluorescence evidence showing that ATRX binds to the short arms of human acrocentric chromosomes where arrays of ribosomal DNA are located (21). ATRX syndrome patients are also characterized by extremely low methylation levels on rDNA, especially in high CpG regions, indicating silent rDNA deficiency (22). However, no further details have been reported other than ATRX and/or DAXX loss have frequently been found in pancreatic neuroendocrine tumors (PanNETs) (23), pediatric glioblastoma multiforme (GBM) (24), low grade gliomas (LGGs) (25,26), and other tumors (27). Thus, the role of ATRX/DAXX/H3.3 in promoting tumorigenesis remains unclear (27). In the present study, massively parallel DNA sequencing data were reexamined in order to trace multilevel epigenetic changes that occur in rDNA during the differentiation of embryonic stem cells (ESCs). Surprisingly, maintenance of silent rDNA was found to require the ATRX/DAXX complex to incorporate H3.3 and establish lysine 9 tri-methylation with the help from the methyltransferase, ESET on P&C regions of silent rDNA. Further experiments demonstrated that loss of the ATRX/DAXX complex converts silent rDNA into upstream binding factor (UBF)-bound active rDNA, thereby releasing transcriptional potency. Coincidentally, LGGs with loss of ATRX exhibit higher ribosomal biogenesis activity, thereby enhancing the more rapid growth of these tumors.

## RESULTS

### 1. Method to selectively subtract background to exclude the noise signal of mapping

Previously, immunofluorescence studies demonstrated that ATRX binds to the short arms of human acrocentric chromosomes where arrays of ribosomal DNA are located (21). Recently reported that Hira-dependent H3.3 is required for ribosomal DNA transcription (28), however whether ATRX/DAXX complex has an epigenetic regulatory role in rDNA remains unclear. Next-generation sequencing (NGS) data has revealed an abundance of epigenetic and protein interaction features of genomes. However, due to the multi-copy repeat and heterogeneity features of rDNA, it is not included in current genome assemblies. Consequently, genomic analyses exclude rDNA. Fortunately, the ability to add one unit of rDNA to currently available genomes has facilitated a re-examination of abundant ChIP-seq data for studies of rDNA (2,29). Since background/noise signals can derive from random protein-DNA or antibody-DNA contacts that are not biologically meaningful or position-specific, it remains a challenge to confirm whether low-level enrichment signal is biologically meaningful. In another case, silent rDNA-specific protein usually shows lesser-level enrichment on rDNA (Fig 2B), just because it is inevitable mistakenly compared with the whole rDNA, not the corresponding silent rDNA as input. Consequently, we determined that selective subtraction of background signal could improve our mapping analysis. According to the signal-noise model (30), “background subtraction” can be performed prior to mapping of rDNA (Fig. 1A). Mapping signals consist of truly enriched fragments and a large number of non-specific fragments which behave as “background”. A crucial parameter of enrichment site detection and error rate control in ChIP-seq data analysis has been defined as normalization factor, *r* (31). The latter allows an input sample to be normalized to a level equivalent to the background noise of a chromatin immunoprecipitation (ChIP) sample. In the present analysis, NCIS (Normalization of ChIP-seq) (32) was used to calculate the value of normalization factor, *r*, thereby making the background ChIP data theoretically equal to *r*\*Input data (Fig. 1A, Step 1). After the background was subtracted, post-processing of data was performed, followed by normalization according to scaling to reads per million (RPM) and division on background-subtracted ChIP and Input data (Fig. 1A, Step 2). We used CCCTC-binding factor CTCF to check this “background subtraction” approach, which localizes upstream of promoters on active rDNA to act as a genome architecture and boundary factor (2). After noise filtered, the unbinding site of CTCF was left a narrow, randomly fluctuating error signal with equally positive and negative values (Fig. 1B).

**Figure 1.**
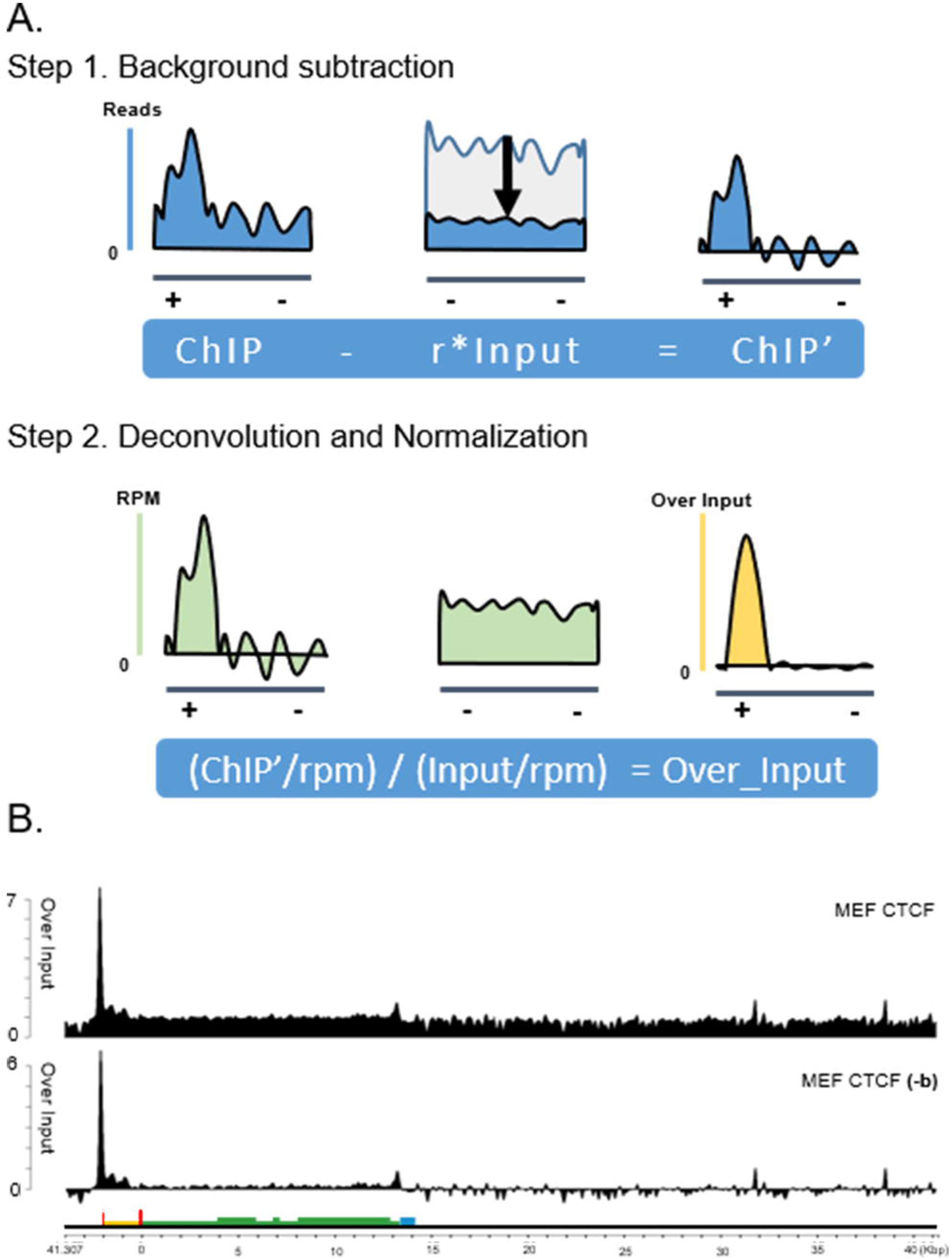
An overview of the method used to subtract background and remove noise signal. A. A simplified overview of the “background subtraction” method. In theory, reads of Input can be scaled equally to background reads of ChIP data according to normalization factor, r. Background associated with the ChIP data can then be deducted by r*Input. Next, background-subtracted ChIP and Input are scaled to RPM and subjected to division for deconvolution. Potential negative (-) and positive (+) sites of enrichment are shown. B. The background signal of CTCF ChIP-seq on rDNA unit is thoroughly removed after performing “background subtraction”, -b indicates background is subtracted.

**Figure 2.**
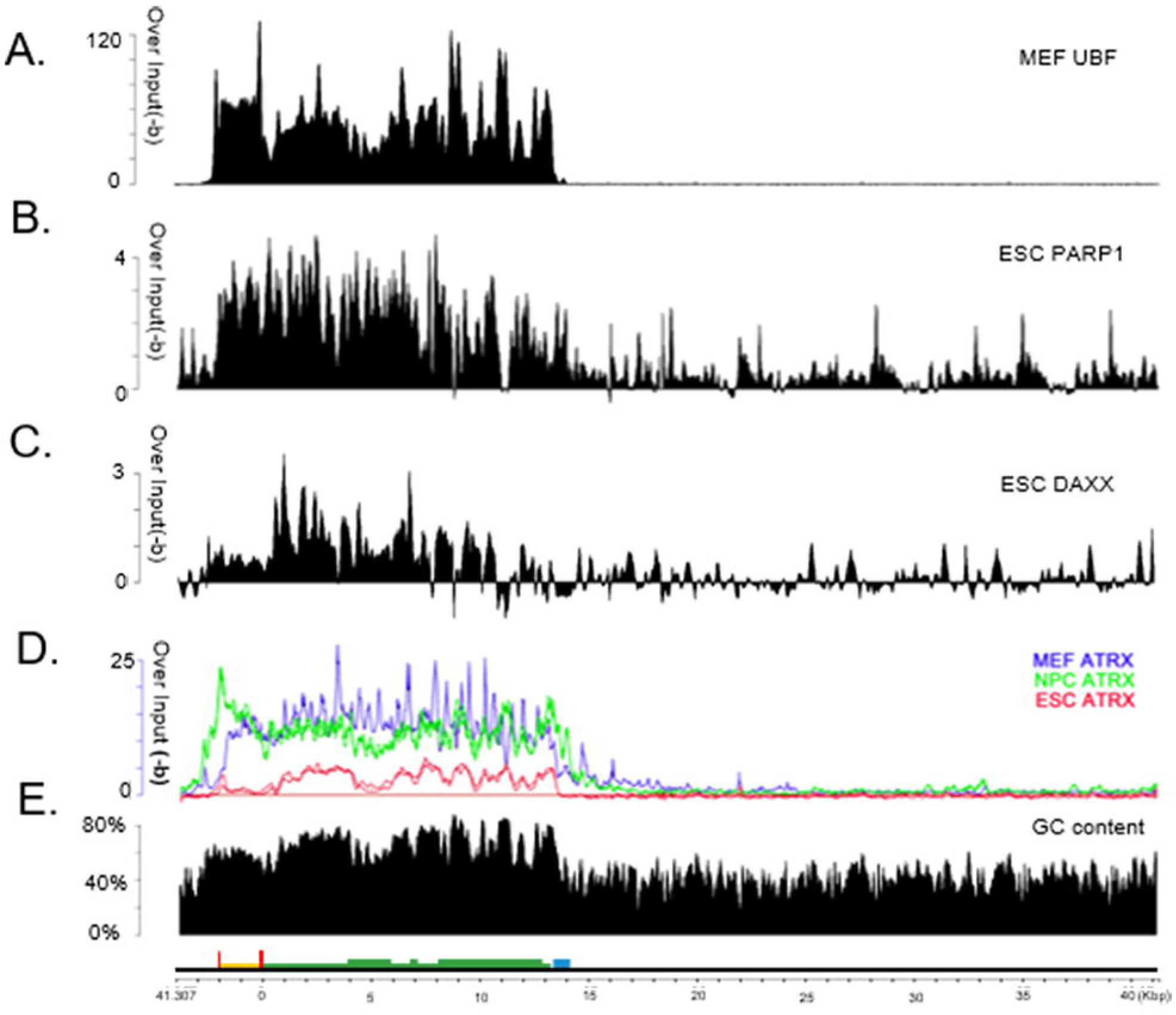
ATRX and DAXX are primarily enriched in the P&C regions of rDNA. A. Maps of UBF on rDNA after background subtraction. B. Maps of PARP1 on rDNA after background subtraction.. C. Maps of DAXX on rDNA after background subtraction. In contrast with the residual error signal in the IGS region, it is more likely that DAXX is located in the P&C region. D. Enrichment of ATRX is observed in the P&C region of rDNA in three different cell types, ESCs, NPCs, and MEFs, after background subtraction (n = 2, 3, and 1, respectively). E. Percentage of GC content across the rDNA sequence examined with a 150-bp sliding window. The P&C region has markedly higher GC content.

### 2. ATRX and DAXX are enriched primarily on P&C regions of rDNA

With this improved method, we firstly reanalyzed the active rDNA associated transcriptional factor UBF. UBF is an essential factor for rDNA transcription that exclusively binds to unmethylated, nucleosome-free regions of active rDNA (2). As it known, UBF specially enriched in P&C regions of rDNA, including spacer and core promoters, enhancers, coding regions, and terminator regions (Fig 2A). Afterwards, we comparatively reanalyzed silent rDNA associated protein, PARP1, which participates in the inheritance and establishment of rDNA heterochromatin (11,33). PARP1 has an actual enrichment primarily on promoter and coding region with reduced level because of its low-proportion of silent rDNA in ESCs (Fig. 2B). As we thought, the enrichment signal of silent rDNA-specific protein is obviously declined, especially in silent rDNA less-existed undifferentiated cell, because it is inevitable mistakenly compared with the whole rDNA, not the corresponding silent rDNA as input. When we study the heterochromatin-related ATRX/DAXX complex, in contrast with the narrow and randomly fluctuating error signal associated with the IGS region, the ATRX/DAXX complex is primarily enriched among P&C regions (Fig 2, C & D). Moreover, ATRX has been found to be consistently localized to P&C regions in ESCs, neuronal progenitor cells (NPCs), and mouse embryonic fibroblasts (MEFs), thereby implying the universality of its association with P&C regions (Fig. 2D).

Considering the previous observations that UBF is closely associated with the GC content of DNA sequences (2) and ATRX also has an affinity for GC content (34), we analyzed the GC content of an rDNA sequence. A distinctly higher level of GC content was found to be distributed in the P&C regions of this rDNA, while the IGS region retained normal GC content (Fig. 2E). The presence of high GC content in the P&C region explains why the ATRX/DAXX complex was enriched in this region (34). However, these observations also suggest a possible competitive relationship between UBF and ATRX when binding toward P&C regions given our current understanding of the functions of UBF and the ATRX/DAXX complex.

### 3. rDNA heterochromatinization happens during cell differentiation

The amount of silent rDNA in cells is believed to markedly increase after cells undergo differentiation, although there are very few bioinformatics studies of silent rDNA to provide evidence for this hypothesis. According to the percentage of methylated rDNA in our assays, we estimated that the percentages of silent rDNA in ESCs and MEFs are 6.76% and 22.57%, respectively (Fig. 3A). Therefore, an approximate 4 times increase in silent rDNA in differentiated MEFs may indicate that a great number of epigenetic events occur in MEFs compared to ESCs.

**Figure 3.**
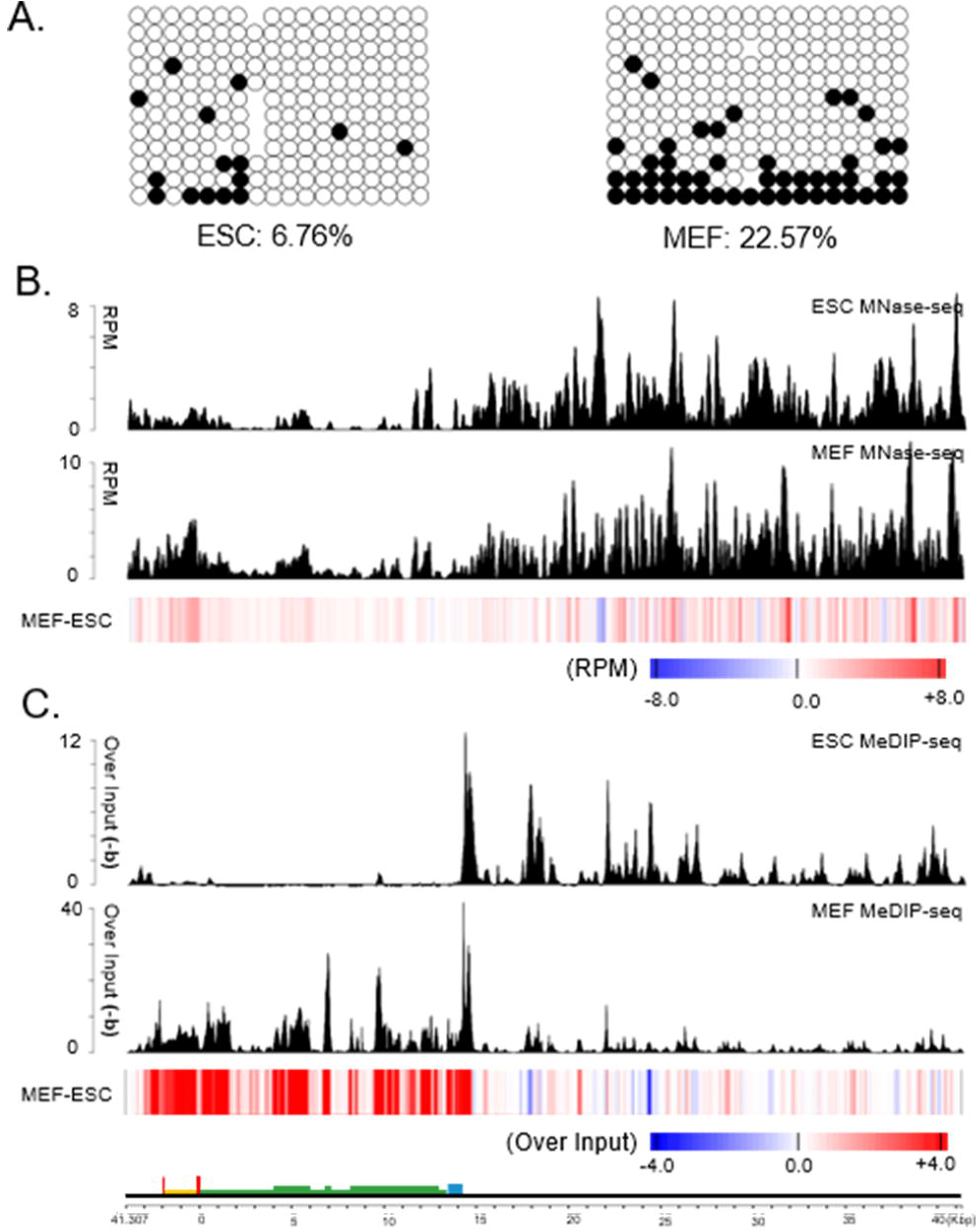
rDNA heterochromatinization happens during cell differentiation. A. Bisulfate sequencing was performed of the 5’ETS (bp 81–507) of the rDNA gene in ESCs and MEFs. Methylated cytosines (black spots) were counted and compared. B. MNase-seq data is analyzed on rDNA from ESC and MEF, and comparison is showed as positive (red) and negative (blue) values below. C. MeDIP-seq data ia analyzed on rDNA from ESC and MEF, and comparison is showed as positive (red) and negative (blue) values below.

To explore the heterochromatinization of rDNA, we compared epigenetic events between ESCs and MEFs with sequence data obtained from the same source. Based on the ability of micrococcal nuclease to digest DNA which is not protected by nucleosomes, we analyzed an undigested portion of nucleosomal DNA (MNase-seq) in order to precisely localize nucleosomes across the genome (35). Markedly fewer nucleosomes were found to occupy the P&C regions compared with the IGS regions in both ESCs and MEFs (Fig. 3B). This result is consistent with previous characterizations of P&C regions of active rDNA as being highly accessible and nucleosome-free (2). In addition, the presence of compacted heterochromatin is supported by the increased histone occupancy observed in MEFs versus ESCs (Fig. 3B, heatmap), thereby suggesting the establishment of heterochromatized silent rDNA during differentiation. The method of immunoprecipitating methylated DNA prior to sequencing (MeDIP-seq) is a technique that is similar to that of ChIP-seq. A methylated 5’ cytosine-specific antibody is used to immunoprecipitate methylated DNA and high-density sites of methylation across an entire genome can be detected (36). Here, we analyzed MeDIP-seq data for rDNA and found a marked increase in DNA methylation in P&C regions than in IGS regions in MEFs compared with ESCs (Fig. 3C, heatmap). These results imply that P&C regions are specifically silenced and heterochromatized by certain mechanism(s) during differentiation. Considering ATRX/DAXX has the same site, P&C regions might be critical site for epigenetic modification of rDNA heterochromanization.

### 4. ATRX/DAXX/SETDB1 drives H3.3K9me3 modifications in P&C regions

Besides, we observed that histone modifications of heterochromatin, such as H3K9me3 and H3K27me3, were widely established on entire rDNA units during differentiation (Fig. 4A), which are consistent with previously published results (6). Moreover, while histone variant, H3.3, has been reported to be related to regions of active transcription via its chaperone protein, HIRA, or to regions of gene silencing via its chaperone, the ATRX/DAXX complex (14), H3.3 displays different changes in its distribution among P&C and IGS regions during differentiation (Fig. 4A). Although it remains unclear why H3.3 modifications are largely absence from IGS regions, we speculate that an increase in H3.3 in P&C regions during differentiation is related to H3K9me3, considering that the chaperone complex, ATRX/DAXX, is specifically enriched in this region (Fig. 2, C & D). With hypothesis verified, ATRX knock-out (KO) and ATRX WT ESCs were compared with respect to H3.3 occupation, showing that only the P&C region was missing H3.3 in the absence of ATRX (Fig. 4B). ChIP-reChIP assays of H3.3 and H3K9me3 followed by sequencing (37) revealed that H3.3 and H3K9me3 were co-enriched exclusively in P&C regions (Fig. 4C), thereby indicating that H3.3 sites are directly modified with tri-methylation of lysine 9 in the tail of histone variant H3.3 to form H3.3K9me3. Correspondingly, when the lysine methyltransferase ESET needed to introduce H3K9me3 is absent, the co-enrichment level of H3.3 and H3K9me3 is markedly decreased in P&C regions (Fig. 4C).

**Figure 4.**
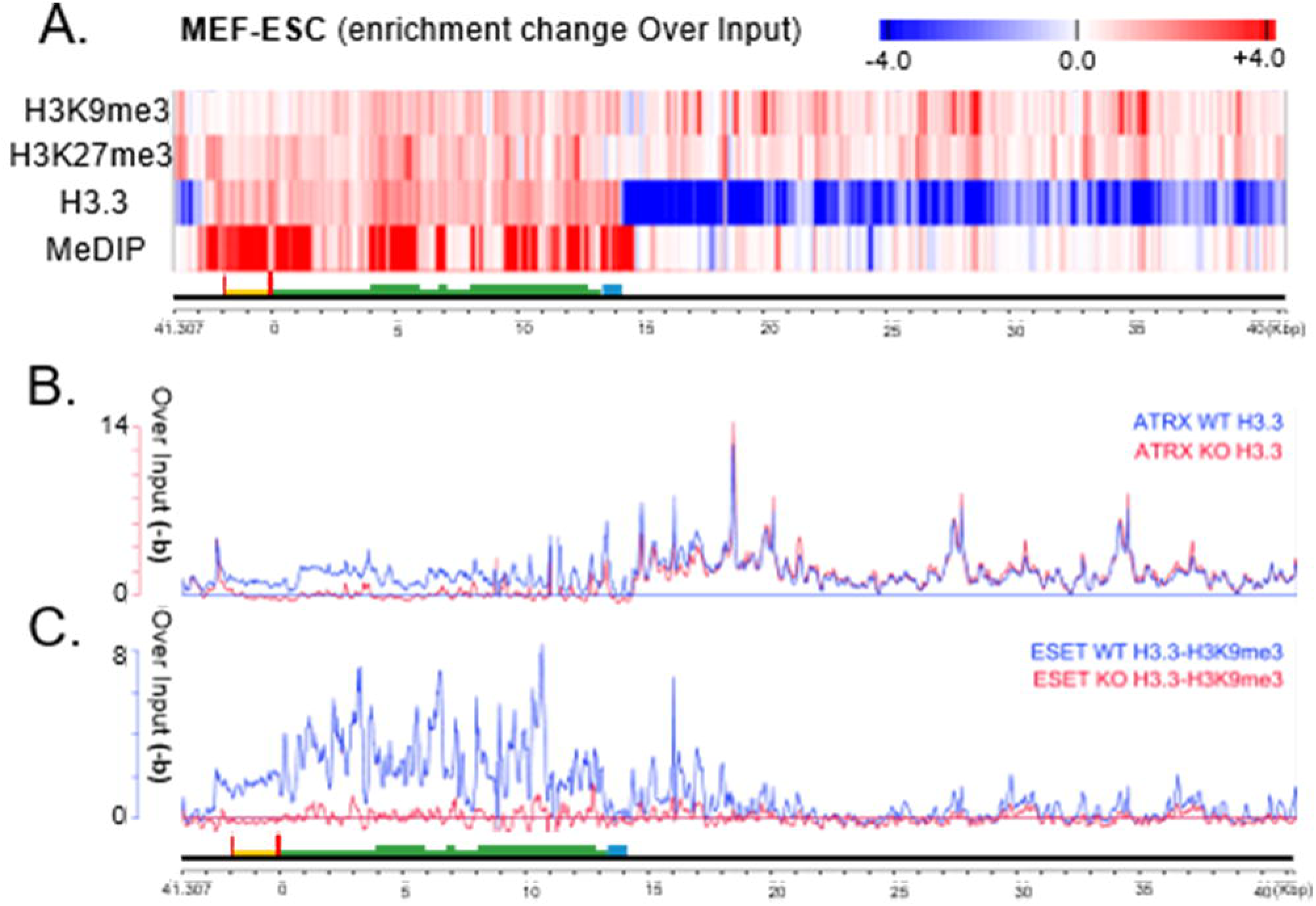
ATRX/DAXX/SETDB1 drives the introduction of H3.3K9me3 modifications in a P&C region.A. Comparison of ChIP-seq or MeDIP-seq data from the same source. The signal (enrichment level over Input with background subtracted) from MEFs was subtracted from the signal in ESCs. Positive (red) and negative (blue) values are shown. B. The H3.3 enrichment level of ATRX-WT and ATRX-KO ESCs with background subtracted. Only the P&C region is markedly reduced in the ATRX KO cells. C. Co-enrichment of H3.3 and H3K9me3 on rDNA in ESET WT and ESET KO ESCs, with background subtracted. H3.3 and H3K9me3 are primarily co-enriched in the P&C region, and this is disrupted when ESET is knocked out.

There are two kinds of lysine methyltransferases, ESET and Suv39h1/2, which catalyze the introduction of K9me3 on H3.3 in cells (37,38). In particular, Suv39h1/2 catalyzes trimethylation of H3K9 at telomeres and pericentric heterochromatin (39,40). However, the mechanism(s) mediating these modifications on rDNA may differ from those used at telomeres and pericentric heterochromatin since no decease in H3K9me3 was observed in MEFs when Suv39h was absent in previous report (41). As mentioned above, KO of ESET markedly abolished the co-enriched level of H3.3 and H3K9me3 (Fig. 4C). In the present analysis, we confirmed that KO of ATRX does not markedly induce a change in rDNA copy (Fig. S1A). Therefore, we hypothesize that the ATRX/DAXX complex is responsible for incorporating H3.3 into the P&C region of silent rDNA and this leads to the introduction of H3.3K9me3 via ESET.

### 5. The ATRX/DAXX complex is required to maintain silent rDNA

To further explore the effect of the ATRX/DAXX complex on rDNA, DAXX was knocked out in MEFs. For these experiments, DAXX^flox/+^ mice were intermated in order to harvest DAXX^flox/flox^ MEFs and DAXX^+/+^ MEFs as controls. KO of DAXX was achieved with virus-mediated Cre expression and 72 h later mRNA and protein levels of DAXX were confirmed to be decreased (Fig. 5A). Next, ChIP-reChIP was performed followed by qPCR to determine the co-enrichment level of H3.3 and H3K9me3 on rDNA. Several primers were designed for various rDNA sites (Fig. 5B). In addition, telomere sites were used as positive controls based on the observation that ATRX/DAXX mediates H3.3K9me3 modifications on telomeres (15,42). In consistence with above ChIP-reChIP-seq data in ESCs (Fig. 4C), H3.3 and H3K9me3 were found to be co-enriched in P&C regions, but not in IGS regions, of rDNA in MEFs (Fig 5C). Moreover, depletion of DAXX resulted in a significant reduction in co-enrichment level of H3.3 and H3K9me3 in telomeres and P&C regions (Fig. 5C), and this result is consistent with the previous observation that mouse ESCs exhibit substantial losses of H3.3 and H3K9me3 at rDNA repeats in the absence of ATRX (43). It has also previously been demonstrated that the methylation level of two special CpG-dinucleotide sites are important for distinguishing active and silent rDNA in mice (44). Thus, we employed the methylation-sensitive restrictive endonuclease, HpaII, to digest unmethylated rDNA and then we performed quantitative PCR to detect methylation levels at -143 bp (Fig. 5D). Bisulfite sequencing was also performed to determine the methylation level of the 5’ ETS region (bps 81–507) (Fig. 5E), which has been identified as one of the primary sites of rDNA methylation in MeDIP-seq data (Fig. 3C). In fact, to a certain extent, the amount of methylated rDNA represents the amount of silent rDNA. Correspondingly, silent rDNA was found to be depleted following the KO of DAXX. This result is consistent with those of a recent study which performed psoralen cross-linking combined with Southern blotting to detect nearly complete loss of silent rDNA in ATRX KO ESCs (43). Therefore, the observations that DAXX depletion leads to a decrease in methylation and H3.3-H3K9me3 enrichment levels suggests that the ATRX/DAXX complex is indispensable for maintenance of silent rDNA.

**Figure 5.**
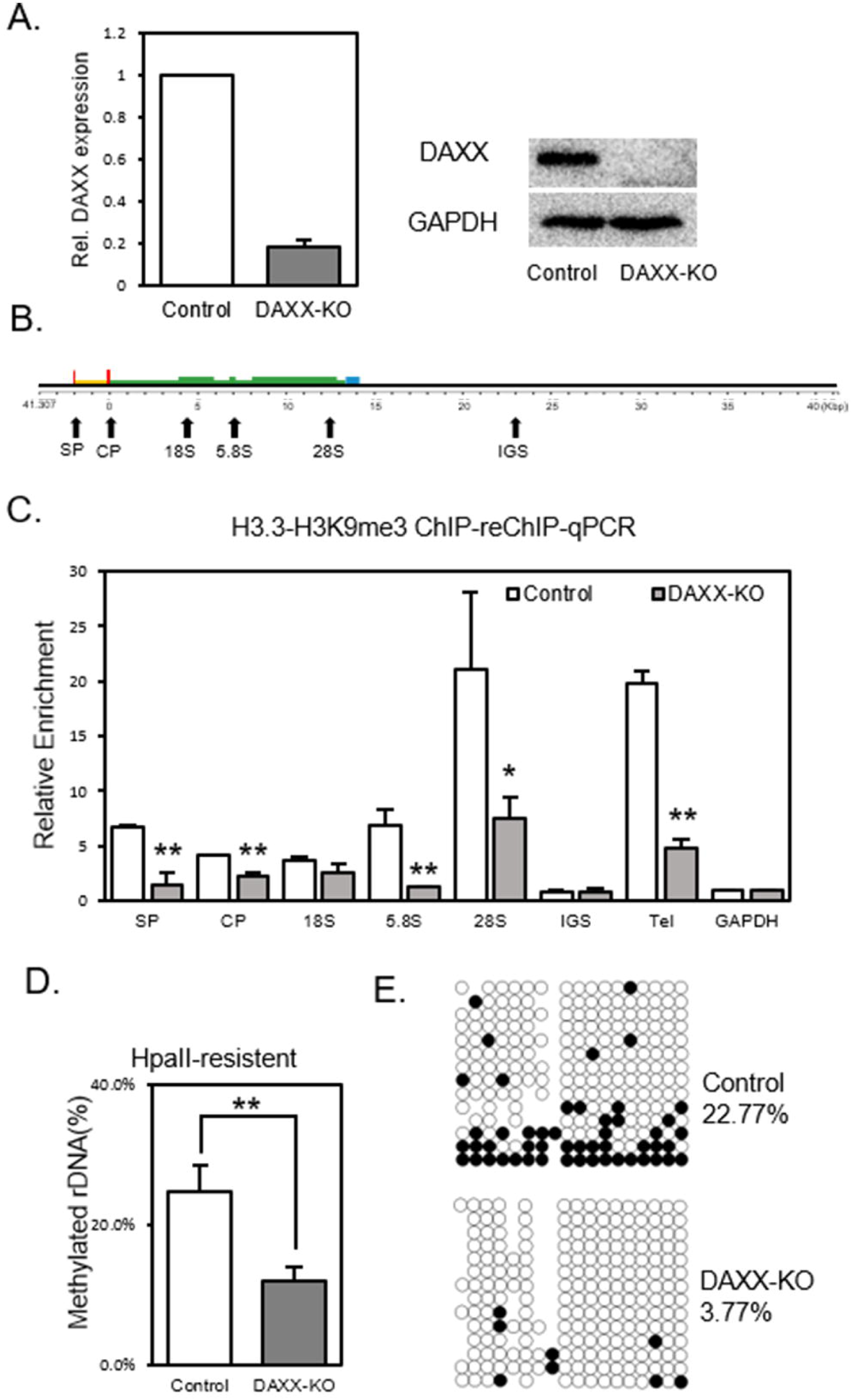
DAXX depletion leads to changed epigenetic modifications of silent rDNA. A. RT-PCR was performed to examine relative *DAXX* mRNA levels (with *GAPDH* used as a reference) in DAXX KO and control MEFs. Western blotting was performed to confirm protein levels of DAXX. B. A diagram of the various primer pairs that were designed for ChIP, ChIP-reChIP-qPCR, the spacer promoter (SP, 43179/43306), core promoter (CP, -105/-1), 18S (4020/4152), 5.8S (6859/6986), 28S (12289/12428), and IGS region (23072/23196). C. ChIP-reChIP followed by qPCR was performed to detect the co-enrichment levels of H3.3 (1st ChIP) and H3K9me3 (2nd ChIP) in DAXX KO and control MEFs. Relative enrichment was calculated relative to GAPDH after normalizing for 5% unbound DNA (see Materials and Methods). Tel, Telomere; served as a positive control. Mean and SD values are derived from three independent experiments. D. HpaII digestion experiments performed in DAXX KO and control MEFs. HpaII-resistant methylated DNA was quantified by primers designed to bind at -143 bp on rDNA. Bars indicate the percentage of methylated silent rDNA in total rDNA, with mean and SD values derived from three independent experiments. E. Bisulfate sequencing of DAXX KO and control MEFs. The 5’ETS (bp 81–507) of the rDNA gene was sequenced. The percentage of methylated cytosines was counted and compared.

### 6. Depletion of the ATRX/DAXX complex releases the transcription potency of rDNA

When silent rDNA is removed, active parts of rDNA have to increase because there is no loss of rDNA copy (Fig. S1B). We performed further experiments to explore this model. We performed silver staining experiments to stain the active regions of rDNA, since elements of the rDNA transcriptional mechanism, including the RNA polymerase I (RPI), UBF, nucleolin, and nuleophosmin B23, exhibit strong argyrophilia (45). In the DAXX KO samples, an increase in the number and area of dark spots, as well as enlargement of nucleolar sizes, were observed (Fig. 6A). These results are consistent with previous silver staining data which demonstrated disruptions in nucleolar architecture and a scatted distribution for dark spots in DNMT1-deficient cells (46). Thus, it has been suggested that active rDNA copies increase in number and surrounding heterochromatin is decondensed in DAXX KO MEFs. Correspondingly, when immunofluorescence spots/areas of UBF binding in DAXX KO MEFs were examined, they were found to have increased in number (Fig. 6B). Additional evidence was provided in a quantitative ChIP analysis of UBF, where DAXX KO MEFs exhibited a significant increase in UBF binding to rDNA compared with its binding profile observed in DAXX WT MEFs (Fig. 6C). These data, as well as data showing a decrease in methylated rDNA (Fig. 5 D & E), indicate that the copy number of euchromatic active rDNA increases upon DAXX depletion. Thus, it appears that the ATRX/DAXX complex is responsible for maintenance of silent rDNA and its depletion prevents the formation of condensed heterochromatic silent rDNA. As a result, silent rDNA is converted into UBF-bound active rDNA (Fig 8).

**Figure 6.**
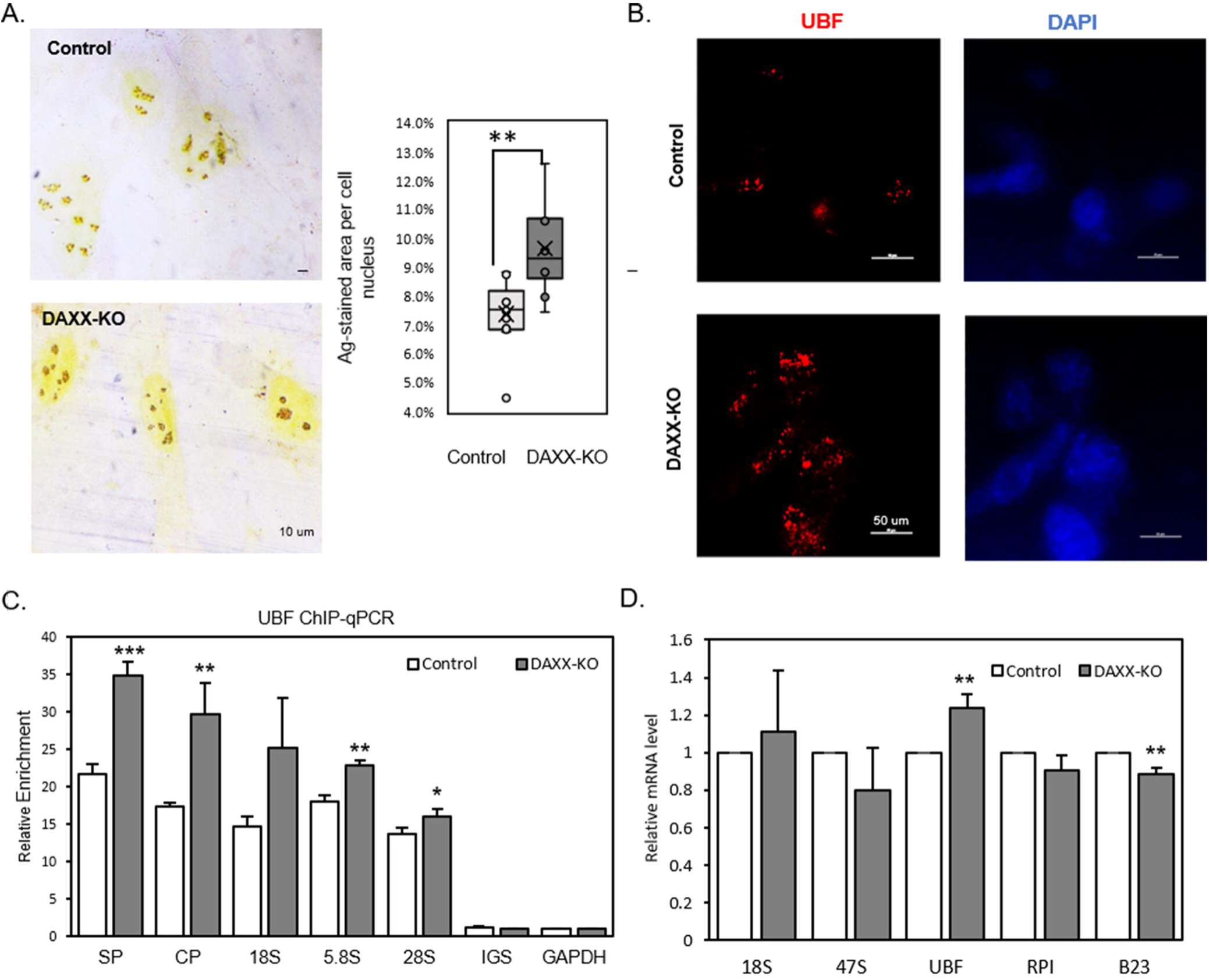
DAXX depletion releases the transcriptional potency of rDNA in MEFs. A. Silver-stained cells exhibited larger scattered dark clusters (representing active rDNA) in the nuclei in DAXX KO MEFs than in control MEFs at interphase. The statistics for ten random cells are shown. Bar = 10 um. B. Immunofluorescence (IF) signals indicate the localization of UBF (red) and DAPI (blue) in DAXX KO MEFs and control MEFs. Bar = 50 um. C. ChIP-qPCR detected UBF-bound rDNA in DAXX KO and control MEFs. Relative enrichment was calculated relative to GAPDH after normalizing for 1% Input. The IGS region, in combination with GAPDH, serves as a negative control. Mean and SD values are derived from three independent experiments. D. Relative expression level of 18S, 47S, UBF, RPI and B23 in DAXX KO and control MEFs. Mean and SD values are derived from three independent experiments.

**Figure 7.**
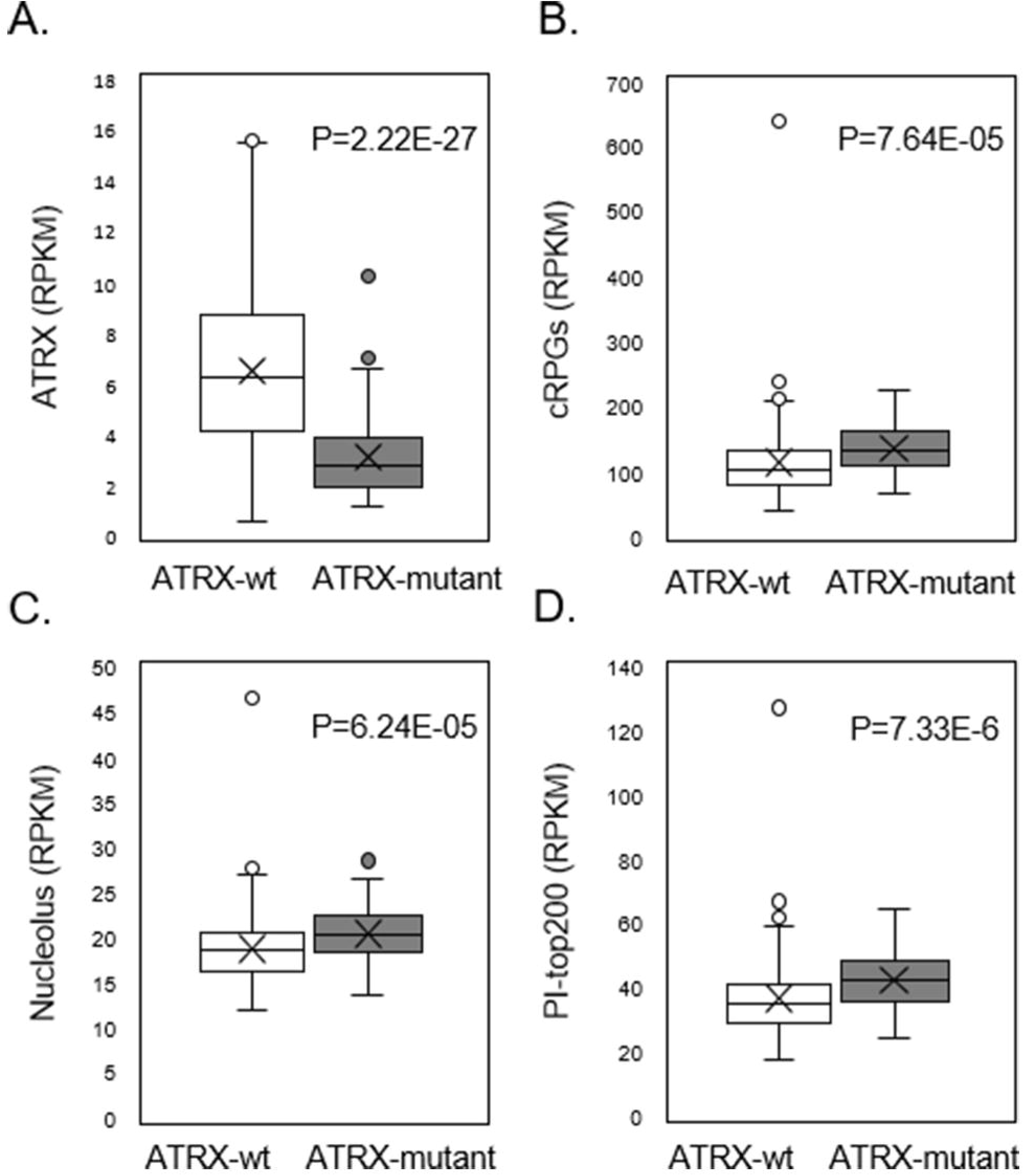
ATRX loss in LGGs promotes ribosomal biosynthesis, nucleolus activity, and cell proliferation. Among the LGG expression data obtained from TCGA, 176 cases and 109 cases carried WT ATRX versus a deficient ATRX due to mutations, respectively. Comparison of the gene expression profiles of these two sets of cases are shown. A. Expression level of ATRX; B. Average expression of cytoplasmic ribosomal protein genes (cRPGs, n = 100); C. Average expression of nucleolus-related genes (n = 799); D. Average expression of the top 200 proliferation-related genes in ATRX-wt and ATRX-deficient LGGs. P-values are shown and were determined with an unpaired two-tailed *t*-test for heteroscedasticity.

**Figure 8.**
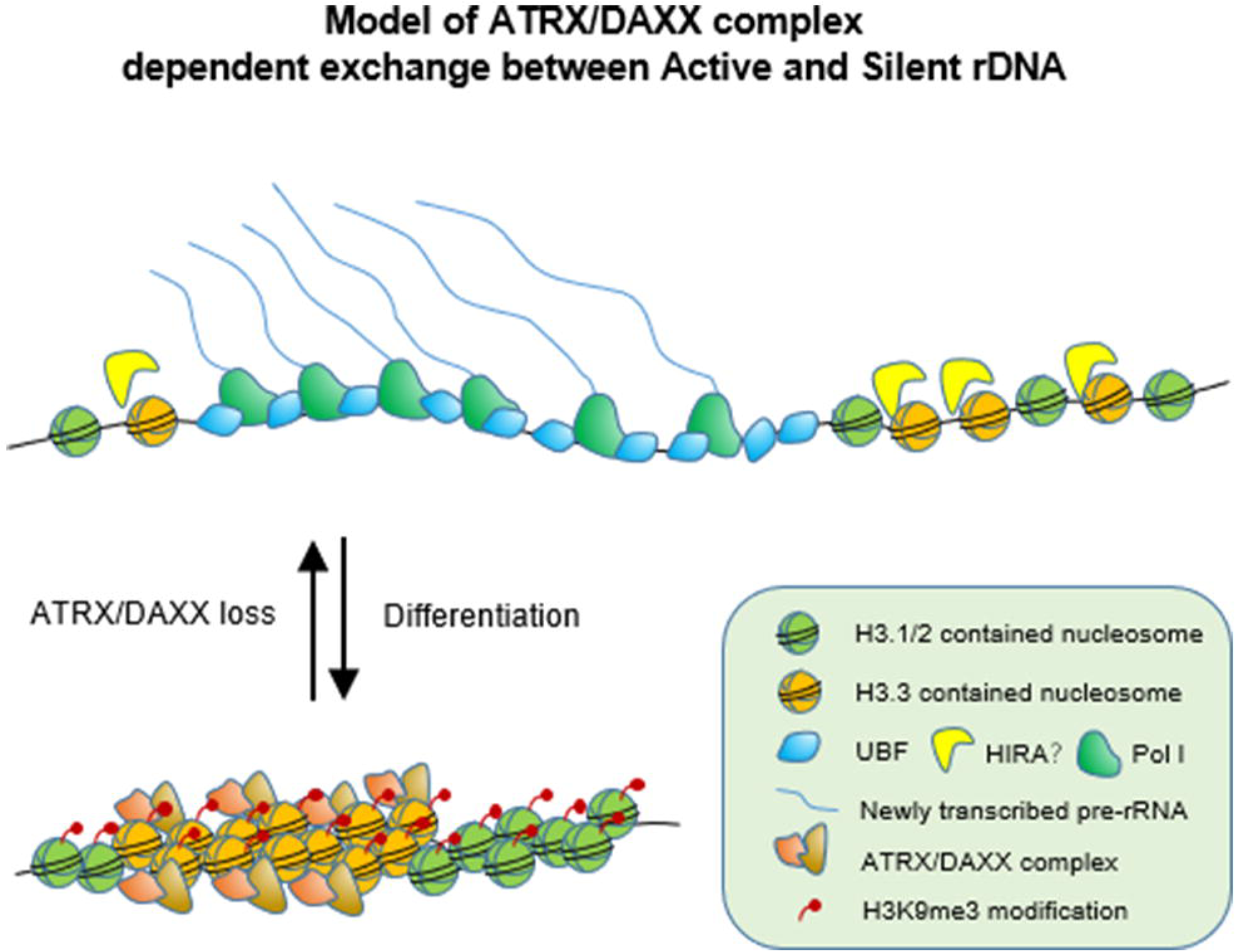
A model showing a proposed transition between active and silent rDNA which is dependent on the ATRX/DAXX complex. In the present study, both UBF-bound active rDNA and ATRX/DAXX-bound silent rDNA were specifically enriched in the P&C region. During differentiation, silent rDNA is generated from active rDNA by the ATRX/DAXX complex, while Setdb1 (ESET) establishes lysine 9 tri-methylation on H3.3. Depletion or loss of the ATRX/DAXX complex result in the conversion of silent rDNA into UBF-bound rDNA and transcriptional potency is released. In particular, LGGs with ATRX loss exhibit higher ribosomal biosynthesis and nucleolus activity, thereby exhibiting more rapid growth.

Considering enlarged ratio of active rDNA, the transcription level would be improved, too. Interestingly, the transcriptional level of rDNA was not augmented (Fig. 6D). Similar results have been reported in DNMT1-deficient HCT116 cells (46) and in PARP1 knock-down MEFs (47). Though transcription factor UBF increases, other transcriptional elements, such as polymerase I and B23 are decreased slightly in expression level (Fig 6D). It is possible that DAXX depletion has a negative effect on cellular physiology and the transcriptional efficiency of polymerase I is downregulated even there is more active rDNA present. However, when knock-down of silent rDNA-related PARP1 does not enhance rDNA transcription in MEFs, it markedly increases transcription during reprogramming (47). Hence, we hypothesize that cells with a higher demand for rDNA transcription may utilize released transcriptional potency.

### 7. ATRX deficiency promotes higher ribosomal biosynthesis activity in LGGs

Cells with a vigorous metabolism usually have higher demands from ribosomal biogenesis, including tumor cells. In fact, the transcription level of rDNA often represents the limit of ribosomal biogenesis (1), and ribosomal biogenesis activity has been strongly linked to tumorigenesis (48). To investigate a possible relationship between ATRX and ribosomal biosynthesis activity, we obtained RNA-seq data for LGGs from The Cancer Genome Atlas (TCGA) Research Network (http://cancergenome.nih.gov/), since mutations in ATRX have frequently been detected in LGGs (49). We obtained 176 cases and 109 cases of *in situ* carcinoma that carried WT ATRX or an ATRX mutant, respectively. We used average cytoplasmic ribosomal protein gene (cRPG) expression to represent ribosomal biogenesis activity as previously described (50), the average expression of nucleolus-related genes to represent nucleolus function activity. Proliferation index (PI) value was determined according to the average expression of 200 proliferation-related genes (51). The LGGs that carried a mutant ATRX had a significantly lower expression level of ATRX compared with the LGGs carrying WT ATRX (Fig. 7A). In addition, significantly higher ribosomal biogenesis activity and higher nucleolus function were detected in the LGGs carrying a mutant ATRX (Fig. 7, B & C). These results imply that release of rDNA transcriptional potency may be fully utilized in tumors that do not express WT ATRX. As a consequence, ATRX-mutant LGGs grow faster with a higher PI index than WT-ATRX LGGs (Fig. 7D). This result is consistent with the observation that ATRX loss promoted tumor growth and reduced median survival in a murine glioblastoma model (52). According to the present study, tumors with loss of ATRX/DAXX may adopt the convenience of rDNA opening, thereby promoting tumorigenesis and oncogenicity (Fig. 8).

## DISCUSSION

Traditional studies of epigenetic mechanisms have focused on gene promoters. Nevertheless, detailed information regarding protein-DNA interactions on whole rDNA are rarely defined. Recently, a numerical deconvolution approach has significantly improved the resolution of protein-DNA interaction maps and has facilitated the construction of sophisticated chromatin structure models for active rDNA (2,53). However, among the available NGS data that derive from ESCs, most contain only a small proportion of silent rDNA. Moreover, existing peak calling routines are used to identify potential factor binding events within a genome (except on rDNA), yet they do not provide sufficient information regarding factor binding at specific sites (54). For the mapping of low-enriched sites, the biggest challenge remains the difficulties associated with estimating where truly enriched sites exist. In ChIP samples, there are a large number of fragments that are generated from non-specific “background” regions throughout the genome (30). Consequently, a crucial parameter of enrichment site detection and error rate control in ChIP-seq data analysis is use of a normalization factor, r (31), which serves to normalize the Input sample to a level equivalent to the background noise of the ChIP sample. Proper estimation of this normalization factor is also important for the identification of weak enrichment sites. There are several algorithms which are able to calculate r, including CCAT (30), NCIS (32), and CisGenome (31). In tools such as MACS (55) and SICER (56), the ratio of the total number of reads in ChIP and Input samples are used to establish a normalization factor (57). Here, we selected NCIS to estimate a normalization factor for eliminating background signal in our maps. As a result, our analysis of silent rDNA epigenetics during differentiation identified a role for the ATRX/DAXX/H3.3 pathway in the maintenance of silent rDNA.

The specific role for ATRX/DAXX identified in the present study involves mediation of the selective co-occupancy of H3.3/H3K9me3 in the P&C region, and not in the IGS region, of the rDNA examined (Fig. 4). Moreover, during differentiation, a greater number of H3K9me3 modifications were present on the entire rDNA, while H3.3 modifications increased in number in the P&C region and decreased in number in the IGS region (Fig. 4A). In contrast, co-enrichment of H3.3 and H3K9me3 was not observed in the IGS region in the bioinformatics analysis performed (Fig. 4C) or in the experiments performed (Fig. 5C). Consequently, we hypothesize that H3.3 in the IGS region correlates with active rDNA independent of H3K9me3. However, it remains to be determined whether this profile of H3.3 is related to HIRA (Fig. 8) and what role it may have.

Previous studies have demonstrated that silent rDNA is established and inherited via the NoRC-pRNA-PARP1 pathway and is also related to the presence of H3K9me2 and H4K20me3 modifications (6,11–13,58). Furthermore, TIP5 (a component of the NoRC complex) and long noncoding RNAs involved in the silencing of rDNA (pRNAs) have been shown to be affected by H3K9me3 modifications in major and minor satellite regions of DNA (6,13). However, no significant changes in H3K9me3 modifications on rDNA have been observed in the absence of TIP5 (13) or during overexpression of pRNA (6). Taken together, these results indicate that a particular mechanism is exclusively responsible for H3K9me3 modifications, and it has not been identified for silent rDNA. In the present study, both the ATRX/DAXX complex and the lysine methyltransferase, ESET, were found to be responsible for H3K9me3 on silent rDNA. For example, in the absence of the ATRX/DAXX complex, silent rDNA was converted into transcription-permissive rDNA. These findings elucidate a supplement to our original mechanism regarding rDNA silencing. However, further study is needed to better understand the relationship between the ATRX/DAXX/H3.3 pathway and the NoRC-pRNA-PARP1 pathway.

Ribosomal biogenesis is strongly linked to tumorigenesis. Consequently, overexpression of rDNA may represent an initiation step during tumorigenesis due to excessive protein synthesis (48,59). Recently, a novel histone H4 variant H4G was found particularly overexpressed in breast cancer tissues and destabilized the nucleosome, which subsequently increased rRNA levels, protein synthesis rates, and cell cycle progression. (60)Correspondingly, recent reports have proposed that epigenetic dysregulation of rDNA may represent a driver of tumorigenesis. For example, when a demethylase of H3K4me3 and H3K36me2, JHDM1B(KDM2B), was knocked down, epigenetic remodeling of rDNA was subsequently induced, and this led to increased invasiveness of transformed and untransformed mammary epithelial cells (61). Meanwhile, depletion of TIP5 has been shown to induce a transforming phenotype in NIH3T3 cells (13). In the present study, it is demonstrated that the ATRX/DAXX complex is indispensable for maintaining silent rDNA, and depletion of the ATRX-DAXX complex converts silent rDNA into UBF-bound active rDNA with fewer methylation sites. Loss of ATRX and/or DAXX has also frequently been observed in PanNETs (23), pediatric GBM (24), in LGGs (25,26), and in other types of tumors (27); and all of these tumors exhibit alternative lengthening in telomeres (ALT) (23,24,26). The latter observation is consistent with the role of the ATRX/DAXX/H3.3 pathway in inhibiting telomere lengthening. However, the role of ATRX/DAXX/H3.3 in promoting tumorigenesis remains unclear (27). In a mouse glioblastoma model, loss of ATRX promoted tumor growth and reduced median survival (52). In clinical PanNETs, low DAXX expression has been found to significantly correlate with a higher Ki-67 index and World Health organization (WHO) grade (62), as well as with a shorter survival period (63). In the present analysis of TCGA data, LGGs with loss of ATRX exhibited higher ribosomal biogenesis activity, enhanced nucleolus activity, and increased proliferative capacity. Thus, a defective ATRX/DAXX complex may promote tumorigenesis and oncogenicity via the conversion of silent rDNA into active rDNA. However, it should also be noted that loss of ATRX/DAXX is not always beneficial to rDNA transcriptional activity. For example, in ATRX-depleted murine ESCs, the ability of the ATRX/DAXX complex to promote rDNA stability was demonstrated, while loss of ATRX or DAXX induced rDNA copy loss and downregulated transcription (43). Copy number loss of 45S rDNA is commonly observed in tumors (64), although it has not been confirmed whether this loss also occurs in GBM and in LGGs. Therefore, further studies are needed to investigate the capacity for the ATRX/DAXX complex to promote tumorigenesis via rDNA.

In conclusion, the findings reported here not only provide a previously undefined mechanism for H3K9me3 modifications of silent rDNA, but they also imply that there may be an association between the ATRX/DAXX/H3.3 pathway and ribosomal biogenesis activity, thereby affecting a key aspect of cell metabolism, growth, and tumorigenesis.

## MATERIAL AND METHODS

### Background Subtraction

To estimate whether low-enriched regions represent a true enrichment signal or a background signal which inherently derives from non-specific signals, we developed a method to effectively reduce background signal. Briefly, according to the signal-noise model (30), *M* and *N* represent the total reads of ChIP and corresponding Input, respectively. We set π_0_ to be the ratio of background signal in *M*. Then, the reads of the background signal were set to π_0_M. As previously described (57), *r* was set as the normalization factor such that:

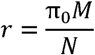

indicating that *r* is the number of times that the background signal is compared with Input. Then, the following formula is applied to obtain the true signal from ChIP data:

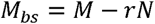

where *M*_*bs*_ represents the ChIP signal with background subtracted. To determine the normalization factor, *r*, NCIS (32) was used to compute its value. Finally, to deconvolute the rDNA (53) and compare it with different cell types, relative enrichment(*M*_*N*_) was calculated over Input. In addition, *M*_*bs*_ and *N* were scaled to RPM, with *RPM* = 1,000,000/total mapped reads:

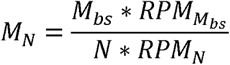

More information regarding the use of python scripts is provided in the Supplementary Materials section.

### Analysis of rDNA in ChIP-seq data

For ChIP-seq and MeDIP-seq, a “background subtraction” step was performed during data processing before a deconvolution protocol was applied as previous described (53). Briefly, raw data were downloaded from the NCBI-GEO database. FastQC was used to check the quality of the data and Trimmomatic was used to cut adapters and low-quality reads for Illumina NGS data. Next, the mouse rDNA repeat (GenBank BK000964.3) was modified so that the origin of the repeat unit was displaced to site 41307 and the pre-rRNA initiation site would be at 4001. Then, the modified rDNA repeat was added to the reference genome version mm10 with the build function of Bowite2. All data were aligned to the newly built reference genome using Bowtie2 with option –k 3. The aligned data were subsequently converted and sorted with SAMtools. Prior to deconvolution, the normalization factor, r, was computed by NCIS. The aligned reads were then extended to 150 bp to calculate the coverage and then they were smoothed by using a 25-bp sliding window. Next, *ChIP* data background was subtracted from *r*Input* as described above. Finally, background-free *ChIP* data and Input data were normalized by scaling to RPM, and relative enrichment (deconvolution) was achieved by dividing the background-free ChIP data by the Input data to remove sequence coverage biases. An available python script is provided in the Supplementary Materials section. In our analysis, region 1974:18089 was designated a P&C region which included spacer and core promoters, an enhancer, a coding region, and a terminator, while the remaining region of the rDNA unit (1:1973 and 18090:45306) was designated an IGS region.

### MNase-seq and ATAC-seq data are shown as RPM

MNase-seq and ATAC-seq have no Input data. Therefore, for the MNase-seq data, we adopted a previously described method (35). The ATAC-seq data was subjected to the same processing method as the ChIP-seq data, although background subtraction and deconvolution were not performed.

### GC content analysis

R package “Biostring” was used to compute GC content in a mouse rDNA sequence obtained from NCBI (GenBank BK000964.3) with a 150-bp sliding-window.

### Antibodies, data, and primer availability

Details regarding the antibodies, data, and primers used are provided in the Supplementary Data section.

### Cell culture and Cre-mediated DAXX KO

Monoallelic DAXX floxed mice were kindly provided by Prof. Fan HengYu (Zhejiang University, China). These mice were cross-mated to obtain DAXX^flox/flox^ fetuses and littermates with DAXX^+/+^ MEFs (WT) used as controls. Primary MEFs were obtained from E13.5 mice and derived MEFs were cultured in high-glucose Dulbecco’s modified Eagle medium (DMEM; Gibco) supplemented with 10% fetal bovine serum. DAXX^fl/fl^ and DAXX WT MEFs were infected with viruses expressing Cre 24 h after inoculation. Then, 72 h after infection, total RNA and protein were extracted for RT-qPCR and Western blotting analyses, respectively. Genomic DNA was extracted for detection of rDNA methylation and copy number.

### Bisulfite sequencing and HpaII-digestion experiments

For bisulfite sequencing, unmethylated cytosines were converted to uracil with an EZ DNA Methylation-Direct(tm) kit (ZYMO Research), according to the manufacturer’s instructions. The 5’-ETS sequence (bp 81–507, GenBankTM BK000964.3) was amplified by PCR, transformed into bacterial clones, and then subjected to sequencing (8). The CpG regions at -143 and -133 were found to be critical for rDNA methylation. In addition, the methylation-sensitive restrictive enzyme, HpaII (R0171S, NEB), was used to distinguish methylated DNA from unmethylated rDNA, as previously described (65).

### ChIP and ChIP-reChIP

Chromatin in two 100-mm dishes of MEFs at 90% confluency were cross-linked with 1% fresh formaldehyde. The chromatin was subsequently sonicated to obtain DNA fragments approximately 500 bp in length. One-tenth of each chromatin sample was subjected to ChIP or ChIP-reChIP assays and 1% chromatin or 5% unbound DNA samples were included in the first ChIP as input or unbound DNA, respectively. Briefly, chromatin was pre-cleared by using pre-absorbed protein A/G magnetic beads (HY-K0202, MCE) according to a previously described protocol (66). Recovered chromatin was then incubated overnight with antibodies at 4 °C, followed by a second incubation with protein A/G magnetic beads for 4 h on a rotating platform. After the protein-DNA complexes were sequentially washed with a low-salt wash buffer (150 mM NaCl, 20 mM Tris-HCl (pH 8.0), 2 mM EDTA, 1% Triton X-100, and 0.1% SDS), a high-salt wash buffer (500 mM NaCl, 20 mM Tris-HCl (pH 8.0), 2 mM EDTA, 1% Triton X-100, and 0.1% SDS), a LiCl wash buffer (250 mM LiCl (746460-100G Sigma-Aldrich), 10 mM Tris-HCl (pH 8.0), 1 mM EDTA, 1% Na-deoxycholate (D6750-10G, Sigma-Aldrich), and 1% NP-40), and twice with TE buffer (10 mM Tris-HCl (pH 8.0), 1 mM EDTA), the protein-DNA complexes were eluted in 1% SDS and 100 mM NaHCO_3_ and then incubated with proteinase K at 65 °C 4 h for ChIP assays. For the ChIP-reChIP assays, the protein-DNA complexes were incubated in 75 ul TE/10 mM DTT (D0632-1G, Sigma-Aldrich) at 37 °C for 30 min, and then a second ChIP protocol was performed. In addition, samples from both assays were digested with proteinase K and DNA was purified before qPCR was performed. Relative enrichment was calculated according to GAPDH after normalizing for 1% Input DNA or 5% unbound DNA.

### Immunofluorescent detection and silver staining experiments

DAXX KO and WT MEFs grown on glass slides were fixed in 4% paraformaldehyde. For silver staining, the fixed cells were initially incubated in a humidity box for at least 30 min before being stained with freshly mixed silver stain medium [2% agar medium with 1% formic acid, 50% AgNO_3_ medium (1:2)] for 10 min. Afterwards, the cells were washed, dried, mounted, and observed with a 100× objective of an oil immersion microscope. For immunofluorescence experiments, the fixed cells were permeabilized with 1% Triton X-100, blocked with 1% bovine serum albumin, then incubated with primary antibodies overnight at 4 °C. The cells were subsequently incubated with appropriate secondary antibodies at room temperature. After 1 h, nuclei were stained with DAPI at room temperature. The slides were subsequently mounted and observed with a microscope.

### Statistical analysis

All experiments were performed in triplicate and data are presented as the mean□±□standard deviation (SD) for each statistical comparison. Two-group comparisons were subjected to Student’s *t*-test and p-values□less than□0.05 (*), 0.01 (**), or 0.001(***) indicated varying degrees of significance for the differences identified.

