## Supplementary material for "The loss of ATRX/DAXX complex disturbs rDNA heterochromatinization and promotes development of glioma": Fig. S1

### Slide 1
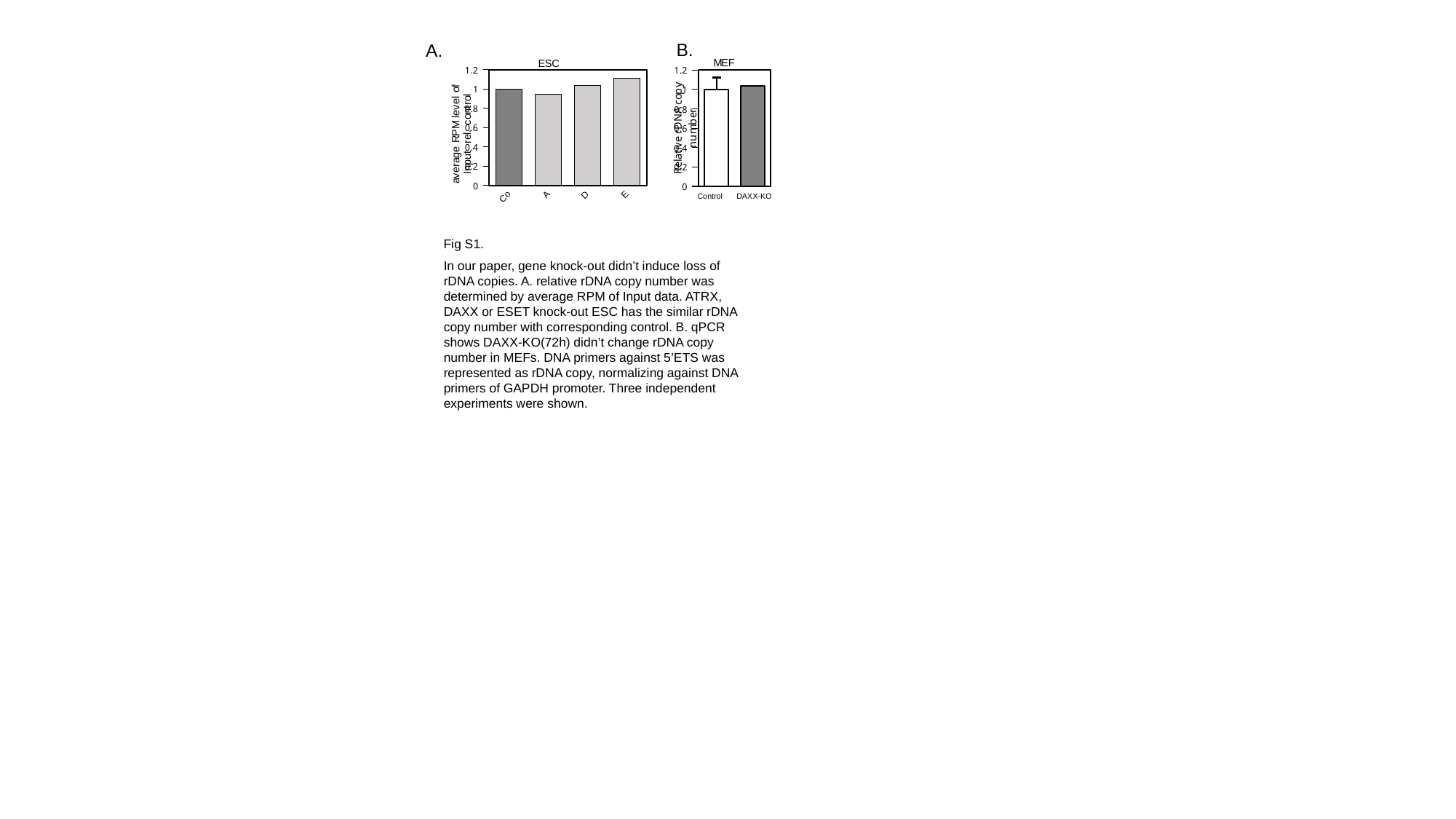

B.
A.
#### Chart: ESC
| Category | |
|---|---|
| Control | 1.0 |
| ATRX-KO | 0.94305986523208 |
| DAXX-KO | 1.0316188844390726 |
| ESET-KO | 1.113667152721276 |
#### Chart: MEF
| Category | |
|---|---|
| Control | 1.0 |
| Daxx-KO | 1.0401455384388614 |Control
DAXX-KO
Fig S1.
In our paper, gene knock-out didn’t induce loss of rDNA copies. A. relative rDNA copy number was determined by average RPM of Input data. ATRX, DAXX or ESET knock-out ESC has the similar rDNA copy number with corresponding control. B. qPCR shows DAXX-KO(72h) didn’t change rDNA copy number in MEFs. DNA primers against 5’ETS was represented as rDNA copy, normalizing against DNA primers of GAPDH promoter. Three independent experiments were shown.
